# Comparison and Characterisation of Mutation Calling from Whole Exome and RNA Sequencing Data for Liver and Muscle Tissue in Lactating Holstein Cows Divergent for Fertility

**DOI:** 10.1101/101733

**Authors:** Bruce Moran, Stephen T. Butler, Christopher J. Creevey

**Affiliations:** Conway Institute of Biomolecular and Biomedical Research, University College Dublin, Belfield, D04 V1W8, Ireland.; Teagasc, Animal and Grassland Research and Innovation Centre, Grange, Meath, C15 PW93, Ireland.; Teagasc, Animal and Grassland Research and Innovation Centre, Moorepark, Cork, P61 C997, Ireland.; Institute of Biological, Environmental and Rural Sciences, Aberystwyth University, SY23 3FL, Wales.

**Keywords:** transcriptome, DNA, SNV calling, bovine

## Abstract

Whole exome sequencing has had low uptake in livestock species, despite allowing accurate analysis of single nucleotide variant (SNV) mutations. Transcriptomic data in the form of RNA sequencing has been generated for many livestock species and also represents a source of mutational information. However, there is little information on the accuracy of using this data for the identification of SNVs. We generated a bovine exome capture design and used it to sequence and call mutations from a lactating dairy cow model genetically divergent for fertility (Fert+, n=8; Fert-, n=8). We compared mutations called from liver and muscle transcriptomes from the same animals. Our exome capture demonstrated 99.1% coverage of the exome design of 56.7MB, whereas transcriptomes covered 55 and 46.5% of the exome, or 24.4 and 20.7MB, in liver and muscle respectively after filtering. We found that specificity of SNVs in the transcriptome data is approximately 75% following basic hard-filtering, and could be increased to above 80% by increasing the minimum threshold of reads covering SNVs, but this effect was negated in more highly covered SNVs. RNA-DNA differences, SNVs found in transcriptome but not exome, were discovered and shown to have significantly increased levels of transition mutations in both tissues. Functional annotation of non-synonymous SNVs specific to the high and low fertility phenotypes identified immune response-related genes, supporting previous work that has identified differential expression in the same genes. Publically available RNAseq data may be analysed in a similar way to further increase the utility of this resource.

**Summary:** The exome and transcriptome both relate to the same protein-coding regions of the genome. There has been sparse research on characterising mutations in RNA and DNA within the same individuals. Here we characterise the similarities in our Holstein dairy cow animal model. We offer practical and biological results indicating that RNA sequencing is a useful proxy of exome sequencing, itself shown to be applicable to this livestock species using a previously untested commercial application. This potentially unlocks public RNA sequencing data for further analysis, also indicating that RNA-DNA differences may associate with transcriptomic divergence.

## Introduction

Whole exome sequencing (WES) is a method used to capture DNA that represents the protein-coding component of a genome. The approach, while only sequencing a small percentage of a vertebrate genome (typically 1-2%), covers those areas where a missense or nonsense mutation would have a higher chance of causing a functional change in the gene product and has most commonly been applied in biomedical studies (Warr *et al.* 2015). Investigating the protein-coding regions by messenger RNA sequencing (RNAseq) has been applied across many vertebrate species and has been widely adopted in livestock species. The specific utility of this kind of research is in the study of gene expression and alternative transcription but there is also potential use in variant calling (McGettigan 2013). In livestock species, WES has been rarely employed, and at the time of writing there were only three bovine exome studies published (Cosart *et al.* 2011; Hirano *et al.* 2013; McClure *et al.* 2014) despite there being an evident utility in causative mutation discovery and in the analysis of ‘extreme phenotypes’ (Emond *et al.* 2012).

The difference in primary technical objectives of WES and RNAseq are quite basic; respectively, the approaches attempt to represent all regions at roughly equal depth and coverage, and represent the level of expression of transcripts. RNAseq inherently results in a dataset of unequal depth and coverage which may also be missing many genes altogether due to not being expressed at the time of sampling. If RNAseq is to be used for mutation analysis, a key outstanding issue to be resolved is determining how WES and RNAseq overlap in their abilities to accurately detect mutations. Several studies using human data have shown that transcriptomic data is a relevant source of information for variant calling and discovery (Cirulli *et al.* 2010; Quinn *et al.* 2013). However, no information is available, neither specifically for bovine nor any other livestock species, as to how to most accurately detect SNV from RNAseq data. Further to this, there has been some controversy surrounding ‘RNA-DNA differences’ (RDD), the proposed result of ‘RNA editing’ (Li *et al.* 2011). Several comments published in response to Li and colleagues indicated that RDD are likely due to issues inherent to RNA sequencing and alignment methods (Pickrell *et al.* 2012; Kleinman and Majewski 2012; Lin *et al.* 2012). With both DNA- and RNA-derived sequence data for the same individuals it could be possible to determine instances of RDD, adding an extra dimension to RNAseq data via WES.

Here we present an analysis of WES of 16 Holstein-Friesian dairy cows. We have previously published RNAseq analysis (Moran *et al.* 2016) which consisted of 96 transcriptomes, six transcriptomes per animal broken into two tissue types (liver, muscle) at three time-points relevant to fertility. Multiple studies have demonstrated that this model of dairy cow fertility has significant differences in reproductive and metabolic performance (Cummins, Lonergan, Evans, Berry, *et al.* 2012; Cummins, Lonergan, Evans and Butler 2012; Cummins, Waters, *et al.* 2012; Butler 2014; Moore, Fair, *et al.* 2014; Moore, Scully, *et al.* 2014). We compared WES and RNAseq derived SNV from this cow model to identify the factors affecting reliability of variant calls in each, especially due to technical differences including uniformity and depth/coverage from each approach. We assessed this independently across the two tissues and also investigated the underlying causes of any RDD observed between the two types of data. We carried out multi-dimensional scaling of the population structure using both WES and RNAseq data, and functionally annotated SNVs from both transcriptome sets.

## Method Overview

### Exome Capture Design and DNA Extraction

Full details of WES data generation, including DNA extraction and exome capture design, are available as Supplementary Material. Briefly, we used the Roche Nimblegen SeqCap EZ Developer kit and to our knowledge this is the first application of the technology to the bovine genome. Genomic DNA was isolated from the same liver samples acquired and used previously in our RNAseq analysis (see Moran et al 2016 for full details of biopsy).

### Bioinformatic Analysis

Raw fastq files were aligned to the UMD3.1 bovine genome, duplicated reads were removed and BAM files were processed using the Genome Analysis ToolKit (GATK v3.1; McKenna *et al.* 2010; DePristo *et al.* 2011) based on ‘best practice’ guidelines (Van der Auwera *et al.* 2013). We then investigated the coverage of exome capture and transcriptome (all exons summed) to determine how well the data represented each. Systematic errors were removed using the Syscall tool (Meacham *et al.* 2011) and all non-biallelic SNV were removed. Hard-filters were then applied to the variant calls, requiring 7 reads supporting the alternate allele, and finally SNV not passing the GATK filters were removed. To determine how these filters impacted the transcriptome data we used absolute expression in the form of fragments per kilobase per million reads (FPKM; Mortazavi *et al.* 2008) to plot expression level versus number of reads.

We used multi-dimensional scaling to visualize proximity of cows and their shared sireages based on the SNV from WES and transcriptomes. Specificity analysis was employed to determine if we could increase the level of true positives (TP, SNVs found in WES and transcriptome) or reduce the level of false positives (FP, SNVs found in transcriptome but not in WES). For those SNV denoted as FP, we classified their occurrence based on several potentially sequencing-related factors. Following removal based on these classifications, we labelled the resulting SNVs as RNA-DNA differences (RDD). Having created TP, false negative (FN, SNV in WES but not transcriptome) and RDD SNV sets, we then investigated the mutational profiles and transition/transversion ratios of each in both tissues and WES, conducting chi-squared tests to determine if a significant difference in transitions/trasversion ratio was apparent. Finally, we annotated the RDD SNV and investigated those non synonymous and found in the differential expression gene set from our previous work for potential functional impact.

## Results and Discussion

### Exome and Transcriptome Design Coverage

Depth of coverage in individual exomes was greater than 10x at 90% of the exome capture design, with all but one sample having 10x coverage over 93% (Figure 1). The percentage of 10x coverage achieved, a standard indicator of quality (Clark *et al.* 2011), was deemed sufficient to proceed with SNV analysis. Coverage across transcriptomes was lower than that of WES (Figure 1), with 10x coverage on average at 85.9% and 83.9% of the expressed regions in liver and muscle respectively. This is likely due to inherently uneven coverage of transcriptomic reads, which yielded very different coverage profiles than WES. For instance, we observed approximately 30% of transcriptome reads in both tissues were aligned to regions with a depth of 100x or more, while only ~1% of the reads from the exome data aligned to such highly covered regions (Figure 1). While SNVs with lower levels of coverage may be missed in transcriptomes due to the reduced regions covered that we report, there is likely to be an increase in support for SNVs in regions that have a higher depth of coverage. Because of this, a strategy increasing the minimum coverage threshold for SNV discovery in RNAseq data may be prudent.

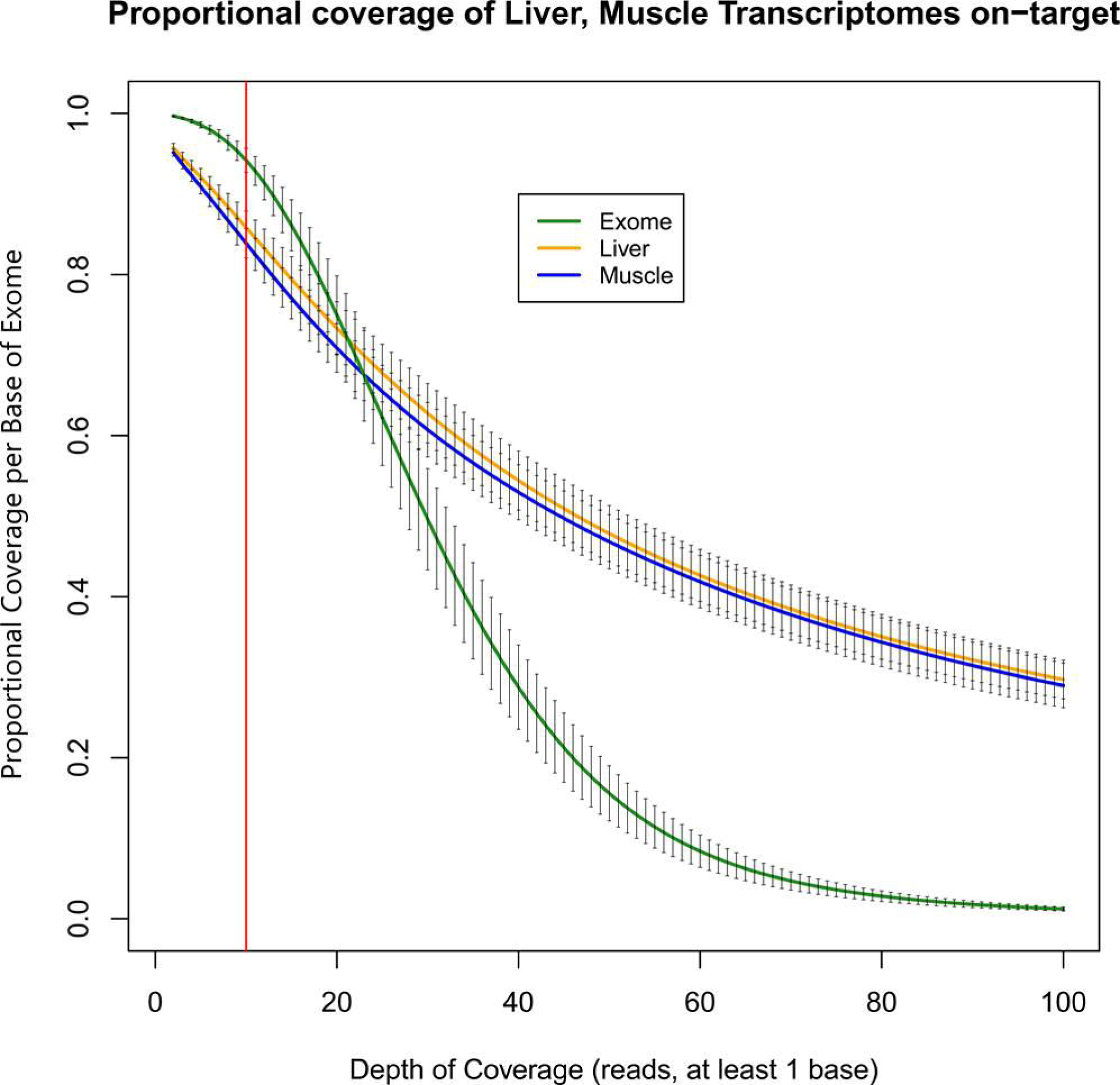

Sequencing metrics are given in Table 1. Per animal, WES had coverage of 97.1 – 97.6% on-target when compared to the exome capture design, and the combined WES data covered 99.1% of the target design. While this difference between individuals is of interest and may simply reflect the random nature of the exome enrichment process, overall less than one percent of the target design could not be captured/sequenced per sample. Increased sequencing levels might potentially be able to ‘fill in’ those regions missed in individual WES based on being found in other individuals if the design was to be used again. The exome capture design can be refined, with more probes added for regions found to be less well covered. The proportion of bases across the exome capture design available for SNV calling for reads passing HaplotypeCaller filtering were 85.4 – 90.6% for individuals and 94.2% for the combined data. This equates to 48,397,629 – 51,344,557 bp in individual exomes, and a total of 53,384,739 bp in the combined data.

Liver and muscle transcriptome designs covered 44,842,138 and 44,506,796 bp of the bovine genome respectively. This equated to 96.8% of the annotated exonic regions of the UMD 3.1 genome annotation in each transcriptome design, and 78.5% of the exome capture design. We therefore concluded that the component of the exome design missing from transcriptomes largely came from the 100bp 5’ UTR of each exon, which amounted to over 22.7MB. The two transcriptomes shared a total of 41,815,289 bp (93.4% of the smaller muscle transcriptome) and combined covered the entirety of annotated protein coding regions. The different regions covered by the two tissues is expected because of tissue-specific gene expression; combining two or more tissues has been suggested as a method to increase the potential for calling variants from transcriptome data (Cirulli *et al.* 2010) and our results show this is likely a good approach. We add that time-point based transcriptomic data could increase the potential of SNV calling from RNAseq data.

A much lower proportion of reads from the transcriptomes passed GATK filters. Output indicated that only 23.9 – 40.8% and 28.9 – 37% of positions, or 10,325,426 – 17,324,585 bp and 10,554,633 – 13,501,253 bp in liver and muscle transcriptomes respectively passed these filters. This eliminated large transcriptomic regions for calling of SNVs. Therefore, the combined transcriptome data which were available for SNV calling represented just 55% and 46.5% of the total liver and muscle transcriptome designs respectively, equating to totals of 24,446,937 and 20,685,045 bp. These transcriptome design sizes were much higher versus the individual ranges, likely due to the transcriptomes coming from three quite distinct time-points at which we might expect different physiological processes to be active, and therefore expression of different genes to be occurring. We tested this hypothesis by combining transcriptomes into time-point specific designs, which showed 84 – 91.1% similarity at each time-point between individuals in liver, and 85.2 – 91.6% similarity in muscle. Again this reinforces our assertion that using RNAseq from multiple time-points is useful for calling SNVs when available.

### Variant Filtering

Using gene expression values to filter RNAseq-derived variants was an obvious strategy. We have found that genes with abundance below an FPKM value of approximately 1, in both tissues, were excluded from the SNV calling procedure because of not passing the HaplotypeCaller filters (Figure 2). This result highlights that genes in regions determined as ‘LOW_COVERAGE’ instead of ‘PASS’ (the threshold being 5 reads coverage) were on average at FPKM < 1, and may be useful for other researchers looking for a baseline threshold for removing genes prior to SNV calling.

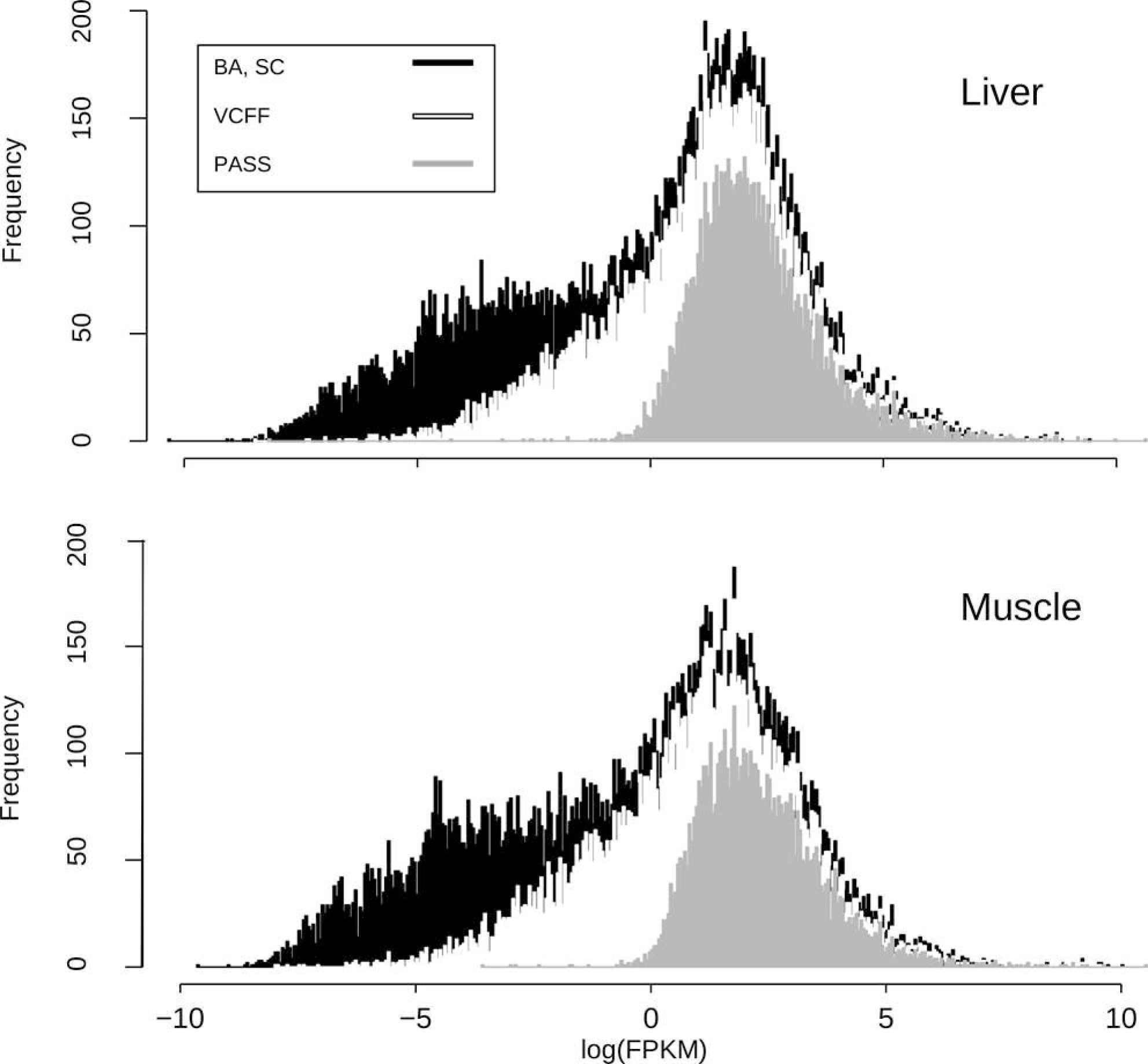

Further filtering of the initial variant calls, beyond the HaplotypeCaller genotyping stage, was carried out by retrospectively removing systematic errors, SNVs with more than a single alternate allele, and SNVs failing our hard-filtering for coverage and occurrence. Initially 219,934 SNVs were called from WES, which was reduced to 135,950 following filtering. This represented a rate of 1 SNV every 417 bp of exome capture design, a rate of 0.2% per base. Of these 88.6% were annotated in dbSNP, the remaining 15,521 being novel. The entire set of SNVs, including those found novel, was used for specificity analysis.

Initially 123,295 and 107,406 SNVs were called from the liver and muscle transcriptomes respectively, which following filtering was reduced to 48,076 and 37,526 SNVs. Of these 82.5% and 82.3% were found in dbSNP, the remaining 8,436 and 6,481 being novel, of which 2,999 and 2,374 (35.5% and 36.6%) were also found in WES. To reduce the possibility of including FP SNVs, technical artefacts of the transcriptome generation process, we only retained those which were also found in WES. This resulted in rates of 1 SNV every 1,043 and 1,334 bp in liver and muscle respectively, rates of 0.095 and 0.075% per base. We also identified 26,067 SNVs common to both the liver and muscle transcriptomes which represented 61.1% and 78.1% of the total SNVs in liver and muscle respectively. Of these 1,795, or 6.7%, were novel.

To determine if WES and transcriptome could accurately identify samples, and those with shared sireage, and MDS plot was constructed using the combined VCF. Insofar as we can visually assess differences between WES and transcriptome derived SNVs, there is some separation (though not complete) between phenotypes, and we have demonstrated that both approaches are reliable for identifying shared sireage (Figure 3). Further, we have shown that this method can act as a quality control step to assure that samples have not been mislabeled, or suffered other handling errors during the laboratory phase of such an experiment.

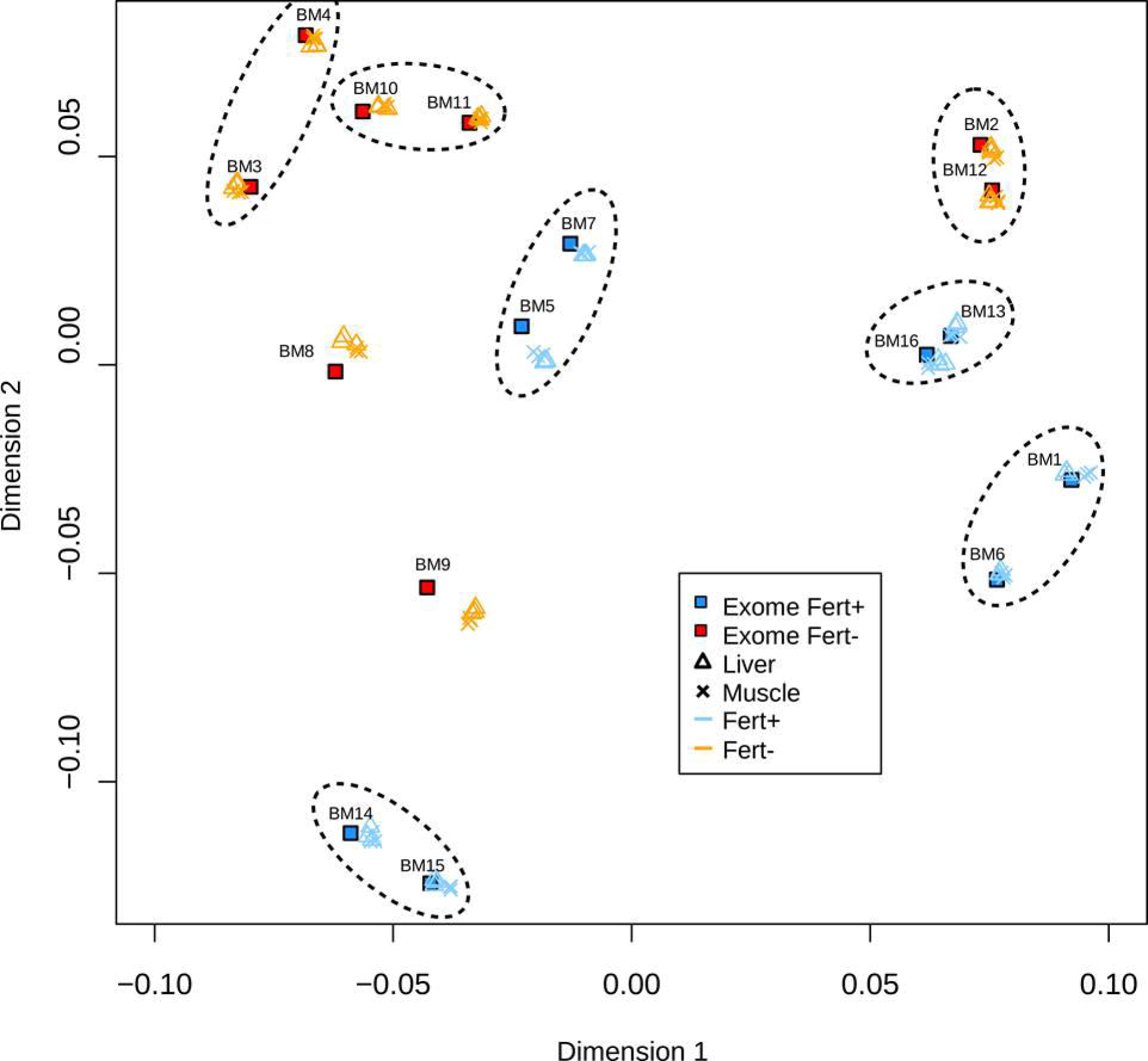

### Specificity Analysis and RNA-DNA Differences

Specificity plateaued at approximately 80% in both transcriptomes, indicating that filtering for depth alone could not improve the FP call rate from RNAseq data (Figure 4). However, increasing the minimum read depth supporting the SNV from 8 to 15 reads in 4 or more samples increased the specificity to almost 85%. However, after this initial increase in specificity at lower coverage we encountered a decline and plateau between 75 – 80% thereafter (Figure 4). We interpret this as resulting from a greater proportion of FP SNVs being called in the most highly expressed genes, and therefore at higher coverage. This may have occurred with a higher number of aligned reads resulting in a greater chance of observing an artefactual mutation. Therefore, whilst GATK uses a maximum allowed coverage for calling mutations, a further strategy that normalises or down-samples the most highly expressed genes in RNAseq data may be prudent prior to variant calling.

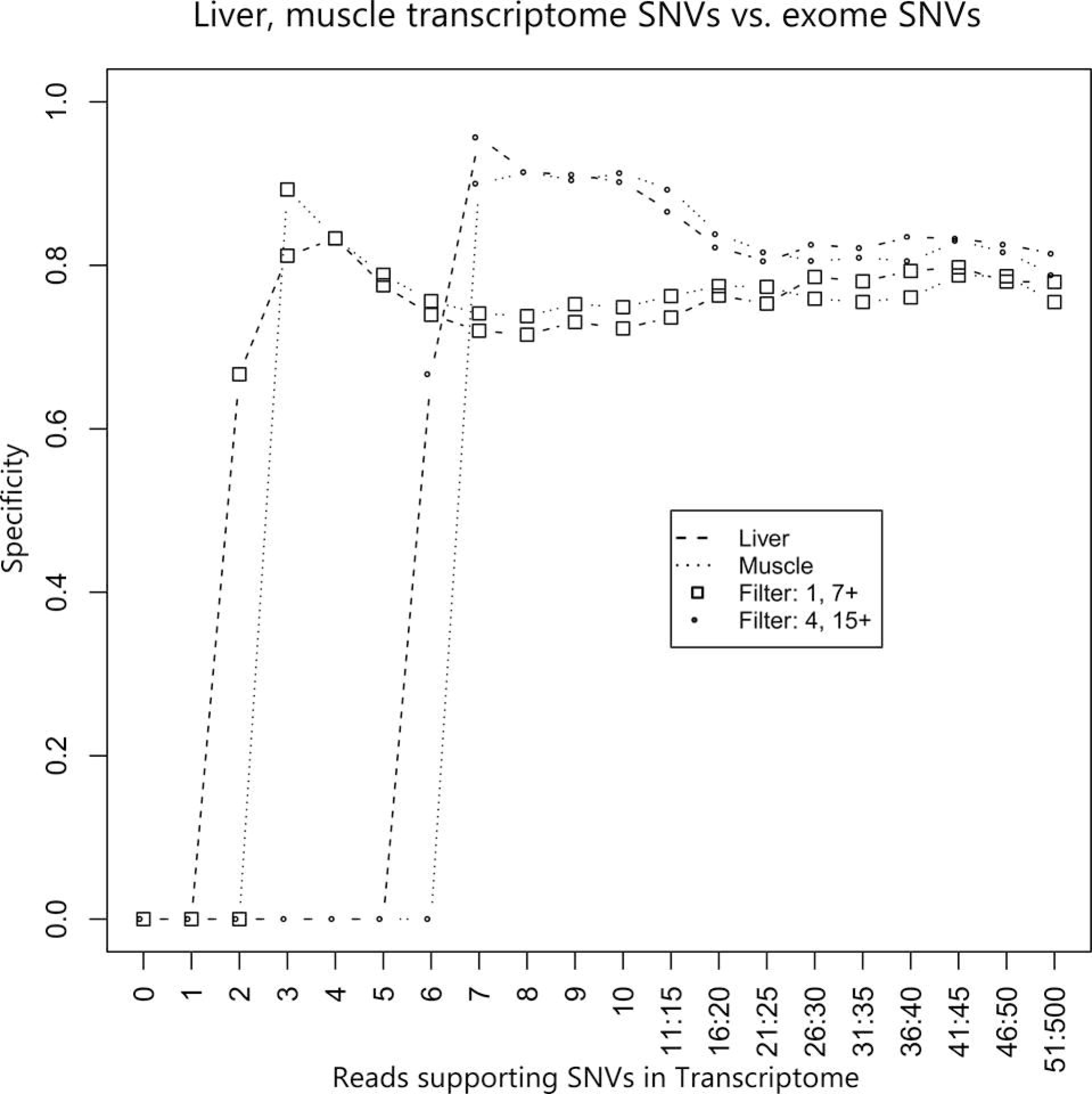

Overall, 9,937 and 7,650 FP SNVs in liver and muscle were also annotated in dbSNP, indicating that they have been independently observed in other studies of cows. We investigated why we had only observed these SNVs in the transcriptomes. Filtering of SNVs that were found in regions of WES with low quality only accounted for 196 and 147 in liver and muscle respectively. This reflected the overall high quality of data generated by WES. Filtering for SNVs that were different to the reference genome and fixed in our population removed a further 175 and 122 FP SNVs. We were also able to account for 408 and 237 FP SNVs found in the last 5 bp of reads; 169 and 90 FP SNVs within 5 bp of splice junctions; and 606 and 333 FP SNVs found within 5 bp of one another in liver and muscle respectively. However, due to overlap between SNVs in each of these categories, totals of 762 and 442 FP SNVs were removed, reducing each set by only 11.8% and 8.5% respectively. This resulted in a final set of RNA-DNA differences (RDDs) of 8,869 and 7,004 FP SNVs in liver and muscle. Of these, 2,160 and 1,936 were unique to liver Fert+ and Fert- phenotypes, and 1,743 and 1,899 unique to muscle Fert+ and Fert- phenotypes, respectively.

The transition/transversion ratio (Ts/Tv) of true positives (SNVs found in WES and transcriptome; TPs) and RDDs for each tissue is shown in Table S1. In WES, Ts/Tv is expected to be ~3 (Marth *et al.* 2011) and we have found a range from 2.63 to 3.57 depending on the SNV set analysed (Table S1). Interestingly, we have found highly significantly different (p < 0.001) Ts/Tv ratio between TPs and RDDs. This manifested as an increase in the frequency of transitions in the RDDs. Furthermore, this result was observed in both liver and muscle indicating a general phenomenon of increased transition mutations in RNA versus DNA.

### Functional Annotation

The numbers of annotated SNVs found uniquely or in both fertility phenotypes in WES and were determined (Figure 5a, b). We have found that of the SNVs unique to a particular phenotype in WES were evenly split, 4,450 and 4,008 were novel, 25 and 26% in Fert+ and Fert- respectively. Of those, 1,158 and 1,031 (again approximately 25% of each set) were found to be non-synonymous (NS). In RDDs we observed lower proportions being NS, with approximately 15% in both liver and muscle in both Fert+ and Fert-. There was low concordance of differential expression (DE) as found previously (Moran et al 2016) with NS SNVs, with totals of 181 (7.7%) and 142 (7%) in Fert+ and Fert-. In liver, there was a lower proportion of NS RDDs unique to Fert+ or Fert- overlapping the DE gene sets than was found in WES, with 15 in each (5% and 6% respectively). In muscle this trend was reversed, and proportionally more NS RDDs in DE genes were found unique to Fert+ or Fert- than in WES, with 22 and 20 (9.1% and 8.4%) respectively. In TPs, there was divergence across these proportions, with 47 each in liver (8.1% and 8.7%), and 38 and 41 in muscle (6.3% and 9.5%) for Fert+ and Fert- respectively. All NS SNVs in liver and muscle RDDs and WES, categorized unique to phenotype, are available in Supplementary Table S2.

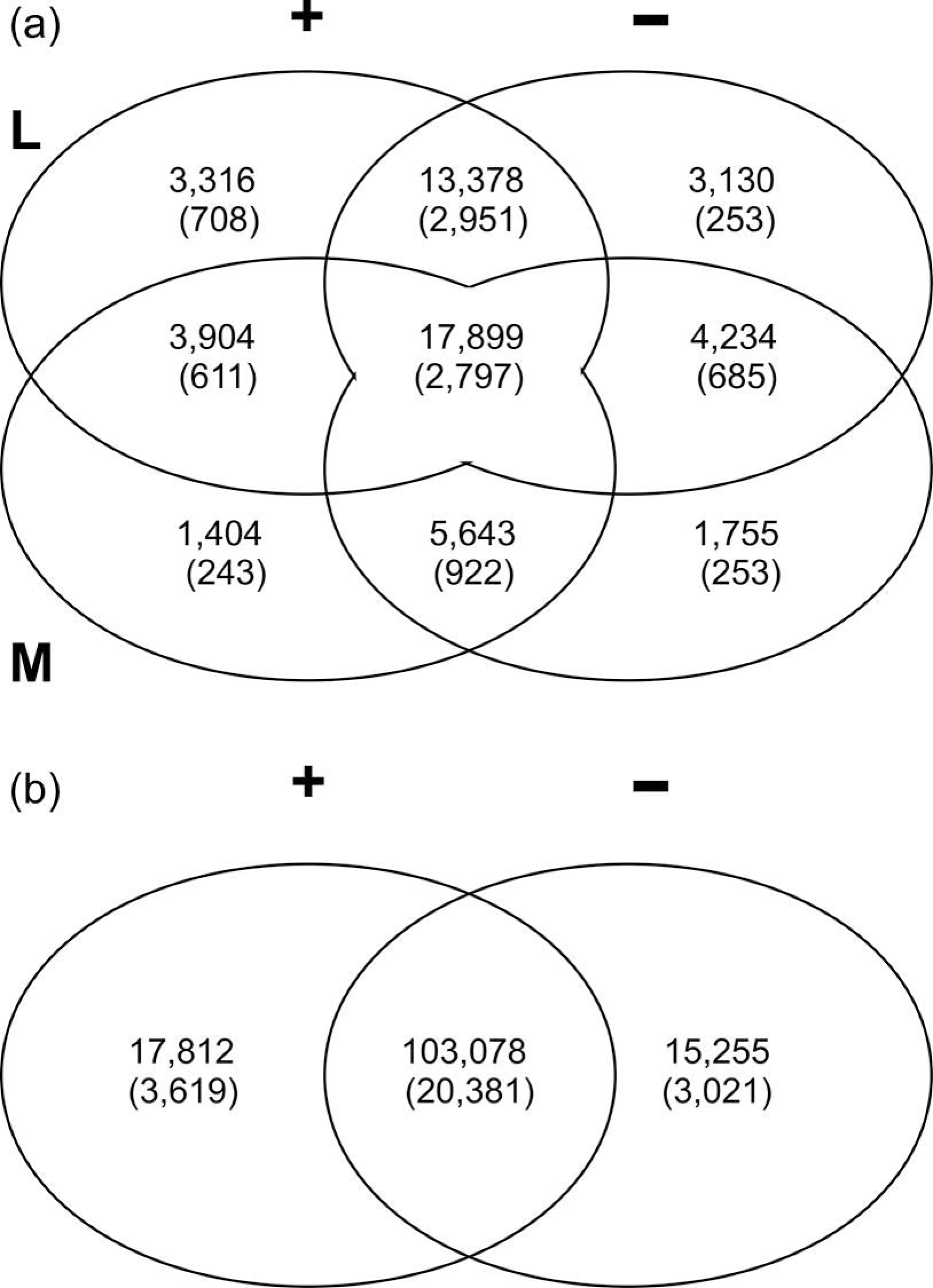

Several of the DE genes with RDDs are of relevance to the phenotype, including the bovine leukocyte antigen (BOLA) super-family of genes. Major histocompatability (MHC) genes have been shown to contain increased levels of mutations, a well-studied phenomenon in humans (Penn *et al.* 2002). Our identification of mutations in BOLA genes found uniquely in RNAseq data raises some interesting questions. In liver, Fert- cows had three unique NS SNVs in *BOLA*, and also one in *BOLA-DRA*, both of which were found to be more highly expressed in Fert- cows in liver in our previous study (Moran et al 2016). Fert+ cows had one *BOLA* and two *BOLA-NC* NS SNVs. Muscle transcriptome showed the same effect, with four NS SNVs in *BOLA* and one in *BOLA-NC* in Fert-; two were found in *BOLA* in Fert+, and one in *BOLA-NC* in Fert+. In WES there were higher numbers, which was expected given the higher quality of the data. *BOLA* contained 8 and 15 NS SNPs in Fert+ and Fert- respectively. Interestingly *BOLA-DQA5* had one NS SNV in Fert-, but none in Fert+, whereas the MHC-related *JSP.1* gene was found with 20 NS SNVs in Fert+ cows, but only one in Fert-. We have found this gene up-regulated at all but one of six time-points. This is an indication of the potential of investigating RDDs to determine their influence on gene expression.

## Conclusions

We have presented the first full characterisation of a bovine whole exome sequencing experiment, with the added benefit of using both a novel exome-enrichment method and complementary transcriptomic data. Previous studies have confirmed the utility of exome sequencing in cows for specific phenotypic and haplotypic studies (Hirano *et al.* 2013; McClure *et al.* 2014; Newman *et al.* 2015). Our findings show that the Roche Nimblegen Developer system has very good potential for generation of WES data based on Ensembl annotation, with very high on-target call rates. Unsurprisingly, we have found that SNV calling in WES data is more informative than using RNAseq data for the same purpose. However, we do not feel that this should exclude use of RNAseq from SNV calling. Indeed, WES exists to both reduce costs of DNA sequencing versus whole genome sequencing, but also because mutations in protein-coding regionsa re expected to be of greater consequence. To this end, RNAseq-called SNVs will inevitably be of value as these genes are by definition expressed. We have shown that some basic steps can be taken to make SNV calling in RNAseq more reliable and we hope these are of interest and might be adopted by others.

Using both data types we have identified a large number of RDDs per transcriptome which significantly deviate in Ts/Tv from our TP SNVs found in both WES and transcriptome. This shows that the results of SNV discovery using RNA sequencing are not identical to WES, and researchers should be aware of this if calling variants without concordant DNA sequencing. However, the ability of transcriptome-defined SNVs to distinguish sireage suggests that taking a conservative approach with SNV analysis resulting from RNAseq can yield biologically relevant and accurate results, and opens up the possibility of mining the vast amount of transcriptomic data available for non-model organisms.

We have also shown that large regions of the transcriptome may not be suitable for variant calling due to low levels of reads passing filters in the gold-standard GATK pipeline. We recommend particular attention be paid to this when using RNAseq data for variant calling. Our key finding that about 20% of SNVs passing all filters were false positives requires further validation. Whilst there is the intriguing prospect of those SNVs being due to processes such as RNA editing or other modifications, there is also the prospect that they are due to GC bias, sequencing errors or because of the evolutionary history of the genes involved. We stress that whilst the finding of these SNVs is an interesting discussion point little functional meaning can be implied without further validation and, even if these RDDs are real, they may be completely benign in terms of gene expression and function.

A stringent approach for identifying SNVs, and particularly the difference in the transition/transversion ratio observed between the TPs and RDDs, is presented. There is an obvious need for further protein and Sanger sequencing work to validate and investigate fully the effect of non-synonymous coding variation found to be divergent between the phenotypes used here. The example of the BOLA gene family, known to have high levels of polymorphism in dairy cows (Hou *et al.* 2012), could be of use in haplotyping animals to further investigate their relationship with fertility. Whilst we have concentrated on NS RDD SNV annotation, we have identified a large number of TP SNVs from RNAseq data. This, along with other public data, represents a wealth of high-quality genetic information that can immediately be harnessed by researchers for a wide variety of studies.

## Methods

### Tissue Biopsy, RNA Extraction and Transcriptome Sequencing Strategy

Tissue biopsies were collected at three time-points relative to parturition (day 0): late pregnancy (LP), day −18 (sd = 7); early lactation (EL), day 2 (sd = 1; EL); and mid-lactation (ML), day 147 (sd = 13). Liver tissue was collected by puncture biopsy as previously described (Cummins, Waters, *et al.* 2012). To collect muscle tissue, a biopsy site on the semitendinosus muscle was shaved and sanitized with 7.5% iodinated povidone and methylated spirits. A subcutaneous injection of lidocaine hydrochloride (2%) was used to anesthetize the area. An incision was made through the skin, and the biopsy instrument (Biopsy Punch 33-37, Miltex GmbH, Riethein-Weilheim, Germany) was used to remove a core of muscle tissue. The incision site was sutured and treated topically with Duphacycline aerosol (3.6% oxytetracycline hydrochloride: Norbrook Laboratories Ltd., Newry, Northern Ireland). Both liver and muscle tissue biopsies were immediately rinsed in saline, blotted dry, snap frozen in liquid nitrogen and stored at −80 °C until RNA extraction.

Total RNA was extracted using a standard Trizol-based method. The tissue sample was weighed and 100 mg cut and homogenized in 3 ml TRI Reagent (Sigma-Aldrich, Dublin) until fully homogenized using a hand-held device. The homogenate was removed to sterile Eppendorf tubes (Eppendorf, UK) and incubated at room temperature (RT) for 5 min; samples were centrifuged at 12,000 × g for 10 min at 4°C, and the supernatant was removed to new sterile tubes. Chloroform was added at 0.2x the volume of homogenate and incubated at RT for 3 min and samples were centrifuged at 12,000 × g for 10 min at 4°C. Isopropanol was added at 0.6 times the volume of supernatant, vortexed and centrifuged at 12,000 × g for 10 min at 4 °C to pellet the RNA. The supernatant was discarded, the pellet was washed twice in 99% ethanol (Sigma-Aldrich, Dublin), and centrifuged at 7,500 × g for 5 min at 4°C. The RNA was re-suspended in 50 μl nuclease-free water (Sigma-Aldrich, Dublin). A kit based protocol (RNeasy Plus; Qiagen, UK) was used to clean the total RNA, removing the fraction below 200 bp and any genomic DNA. RNA quality was assessed using the Bioanalyser 2100 (Agilent Technologies, UK) with the RNA Nano chip.

Library preparation was conducted using the Truseq v2 kit (Illumina, UK) following the protocol. The Bioanalyser 2100 (Agilent Technologies, UK) was used to visually determine quality of the libraries with a DNA Nano chip. Concentration was determined using the KAPA Library Quantification (KAPA Biosystems, USA) qPCR method to allow equimolar pooling of each library. Four pools of 24 libraries were made, the maximum possible based on the number of barcoded adapters available at the time. Each pool was sequenced for 100 bp using a paired-end strategy across 3 separate flowcell lanes on an Illumina HiSeq 2000 (Illumina, USA) to minimize technical variation, resulting in 12 lanes of HiSeq data. RNAseq data are available in the Gene Expression Omnibus (GEO) at accession ID GSE62159.

### Exome Design, DNA Extraction and Sequencing Strategy

We used the Roche Nimblegen SeqCap EZ Developer kit to perform the experiment and this was, to our knowledge, the first study using this method on the bovine genome. This method requires a user-generated exome sequence from which probes are made; we used the exome regions from the UMD.3.1 Ensembl 70 release genome transfer file (GTF) including 100bp of 5’ sequence of all exonic regions of which there are 227,647. Our exome target design covered 202,899 consolidated regions over 56,671,697 bp. Probes were allowed to have up to 5 total matches in the genome and 5 mismatches within those, although 92.5% of probes were unique to a single genomic position and a further 4.5% had only two possible matches within the genome. Probe length was 200bp with approximately 2 million probes covering the exome. Probes targeting mitochondrial DNA were at 1/5^th^ the concentration of nuclear DNA.

Genomic DNA (gDNA) was extracted from liver and muscle tissue biopsies using the Zymo Quick-gDNA^TM^ MiniPrep kit (Zymo Research Europe, Germany). We processed samples in triplicate as per the protocol with starting volume of 25mg of tissue homogenised with a Precellys bead beater in 500ul of Genomic Lysis Buffer. We eluted in 50ul of DNA Elution Buffer and proceeded to the Zymo Genomic DNA Clean and Concentrator^TM^ to clean and concentrate the gDNA. We used an input of 25ul and output of 10ul. This gDNA was sent to a core facility, which conducted the library preparation, hybridisation with exome capture kit and sequencing on 3 lanes of HiSeq 2000. Our strategy for hybridisation and sequencing was to make one single equimolar pool of all 16 gDNA libraries, each having different adapter barcodes to allow us to de-multiplex sequence data into the correct sample for analysis. We then split the pool in four parts and performed hybridisation captures on each part using 2x capture reagents, resulting in 1x capture for every 2 samples. All captures were then re-pooled and then split into three parts for sequencing on three HiSeq lanes. We used pooling strategies to attempt to reduce variation in hybridisation, and because we were ultimately pooling the captured libraries to sequence over three lanes. Thus, the first pooling of library ensured the samples were well mixed prior to capture, and then the capture was well mixed to give as even a distribution of sample to lanes, and previously sample to capture reagents, as possible. Exome data are available in the Gene Expression Omnibus (GEO) at accession ID 62160.

### Sequence Alignment

WES data were aligned with BWA using the ‘sampe’ algorithm (Li and Durbin 2009). Resulting SAM files were sorted, duplicates removed and read groups added using Picard Tools (available from: www.picard.sourceforge.net). Pre- and post-duplicate removal statistics were generated for SAM files using Samtools Flagstat (Li *et al.* 2009) to determine numbers of reads properly paired and aligned. SAM files were converted to BAM files using Samtools View which were subsequently used for variant calling.

Transcriptome data were aligned using STAR aligner (Dobin *et al.* 2012) with a ‘two-pass’ strategy for novel splice junction annotation. The aligner was first run using the same gene transfer format (GTF) file that was used in designing the exome from which canonical splice junction boundaries could be determined. This preliminary run resulted in a splice junction database per sample; we collected all such splice junction databases, removed junctions which had less than 20 occurrences, and used that set of junctions for our ‘second-pass’ alignment, instead of the GTF file. We then sorted, removed duplicates and added read groups with Picard Tools, generating pre- and post-duplicate removed SAM file statistics using Samtools Flagstat. SAM files were once again converted to BAM files using Samtools View and we proceeded to variant calling.

### Variant Calling

The Genome Analysis Tool Kit (GATK) version 3.1 (McKenna *et al.* 2010; DePristo *et al.* 2011; Van der Auwera *et al.* 2013) ‘walkers’ (algorithms for a variety of data processing) were used under their ‘best practices’ guidelines for sequence variant calling. For transcriptome BAM files the SplitNCigarReads walker was used to hard clip bases following ‘N’ in the CIGAR string. This removes all “overhangs” into the intronic regions. Having previously defined the splice junctions in STAR we were confident that this approach did not result in elevated sequence removal compared to the overall greater alignment and ultimately potentially greater variant calling accuracy. This walker also reassigns mapping qualities to change MAPQ 255 from STAR, meaning the read aligned uniquely (i.e. to only one position of the genome). The 255 score is determined as ‘unknown’ by GATK walkers, and so all 255 scores were changed to 60 which is used downstream in GATK to mean uniquely aligned reads.

WES and RNAseq were then processed almost identically. The RealignerTargetCreator and IndelRealigner walkers were run on the BAM file with the Ensembl 74 version dbSNP variants given as known variants. The BaseRecalibrator walker was then used to recalibrate scores around those known variants and PrintReads walker was used to generate a recalibrated BAM file. The HaplotypeCaller walker was used with Phred-scaled emit and call confidences of 30 for WES and 20 for RNAseq in the ‘GVCF’ mode and with the BED file of our exome target design to guide calling. This resulted in a single genomic variant call format (GVCF) file per sample. The entire set of GVCF files for each of WES, and the liver and muscle transcriptomes separately, were then ‘joint genotyped’ using the GenotypeGVCFs walker. This generated a single joint genotyped GVCF file containing information for all 16 exomes at every variant site found in any of the exomes, and 48 samples each in liver and muscle transcriptomes, at each variant site found in that transcriptome in any of the 48 samples. We call the joint GVCF files the ‘variant call files’ (VCF) henceforth for clarity. We removed all but the SNVs in which a single alternative allele was found in the variant call file. This is because the ENSEMBL version 74 bovine dbSNP file contained ~21.2M positions of which ~19.7M SNPs (~93%) have a single alternate allele and so other variants might be under-represented, making them more difficult to accurately call. We retained a distinction between the two transcriptomes (i.e. liver and muscle tissue) because this might inherently be of interest: (i) to independently validate results; (ii) for functional annotation of SNPs.

### Systematic Error Removal

To remove possible systematic sequencing errors we used the Syscall tool (Meacham *et al.* 2011). For WES and each transcriptome separately, we took each realigned, recalibrated BAM file (the file from which SNP calls were made), and also took the set of all heterozygous SNVs from the concordant GVCF file per sample. To save computational time we ran Syscall over the first sample, then took all systematic errors as called by Syscall, removed them from the second samples heterozygous SNVs and proceeded through all samples. A pooled error-file was produced and any SNV positions in that were removed from the VCF. This approach of analysing the BAM files individually in turn was completed in approximately a third of the time estimated for carrying out the same analysis on a single BAM file containing all samples (factoring in the time taken for merging of all exome BAM files, sorting and indexing).

### Hard-filtering of SNVs

We had no genotype data from other sources (e.g. whole genome sequence or SNV array) for the animals used in this study and so it was not possible to conduct variant quality score recalibration (VQSR) in GATK. We therefore conducted a series of filtering steps on our variant call files to retain data in which we had higher confidence. We required each SNV to have at least one alternate allele in at least one sample with greater than 7 reads supporting the alternate allele. If the sample was heterozygous the alternate allele also had to be represented by at least 25% of the reads covering the position as well as having at least 7 reads supporting the alternate allele. To test if filtering affected the specificity analysis we increased the hard-filtering thresholds, requiring the SNV to be found in at least 4 different samples, and at depth of at least 15 reads.

### Reads Passing GATK Filters

To determine what proportion of WES or transcriptome could be used in SNV calling by the HaplotypeCaller we ran DiagnoseTargets. This walker applies the same filters to the BAM as HaplotypeCaller and outputs a file with intervals tagged as ‘PASS’ or other indicators of filtering (‘NO_READS’, ‘LOW_COVERAGE’), where ‘PASS’ represents intervals which pass all the filters for quality which were applied. We merged the intervals not passing filters into BED format and took total bases divided by total size of the exome design to get a percentage of the total target design in which SNVs were not callable. Because transcriptomes were inherently smaller than the entire exome design we used two transcriptome designs, as well as the exome design as above, to get percentage of callable regions per each transcriptome.

### Coverage Analysis

We used Bedtools genomecov (Quinlan and Hall 2010) to determine coverage across the genome of each of the realigned, recalibrated BAM files, with output in BED format. BED files contained a chromosomal interval between which reads in the BAM were mapped, as well as holding information on the number of reads supporting that interval. Bedtools intersect was used to determine intervals that were common between individual BED files. For exomic BED files, we intersected the individual BED files with our exome design to determine how well the exome capture process had worked, i.e. how much of the entire exome was covered by at least one sequencing read. We also used transcriptome BED files to determine how well transcriptome intersected the exome design. We combined all WES and transcriptome BED files, merged using Bedtools merge to join overlapping regions, and intersected this with the exome design to show what intervals, if any, were missed entirely by WES, and what regions of exome were not covered by the transcriptomes. We used each merged transcriptome separately to calculate percentage of callable regions in each.

We expected expression of different genes at the different time-points that were represented in the transcriptomes, but did not expect all genes to be expressed in either tissue. Therefore, we used the absolute abundance of genes in fragments per kilobase per million reads (FPKM; Mortazavi *et al.* 2008), and investigated in which genes variants could be called based on passing filters. This gave a distribution of FPKM prior to and following filtering using the preliminary ‘single alternate allele’ and removal of systematic biases, after our hard-filtering, and after the HaplotypeCaller ‘PASS’ filtering.

We also used BED files coverage information to determine the proportion of coverage at depth. We summed all depths (giving the total number of reads covering the transcriptome) and then sequentially took the reads at the next depth minus the reads at the last depth from the total to show what proportion of reads was left at that particular depth. For example for each depth within the window 1 - 10X having 5 occurrences in the BED we have 50 total reads (proportion is 50/50 = 1). Taking away depth=1x leaves 45 reads at depth=2x (proportion of 45/50 = 0.9) and so on.

### Multi-Dimensional Scaling Plot of Combined Exome and Transcriptomes

We created a combined VCF containing all SNV data following filtering from WES and both transcriptomes. Where an SNV was found in only one or two datasets, the remaining set(s) had missing values inserted. Because the exome and two transcriptome sets came from the same animals, we visualised the identity-by-state of the animals with the combined VCF. We used PLINK 1.9 (PLINK v1.90b2a) multi-dimensional scaling (MDS) from the cluster method. We plotted the first two dimensions using R statistical software (R Development Core Team 2012) and as all cows except two had shared sireage with one other cow in the study, we superimposed this information onto the plot.

### Specificity and Transition/Transversion Analysis

We determined what overlap occurred between WES and transcriptomes by specificity analysis. In this context, we defined specificity as the proportion of all SNVs detected that are true positives (Cirulli *et al.* 2010). For this purpose SNVs were categorised as true positive (TP) when the SNV was found in both WES and transcriptome VCFs; false positive (FP) when an SNV was found in transcriptome but not in WES; false negative (FN) when an SNV was found in WES and not in transcriptome. We used an in-house Perl script available at from the authors on request) to parse the WES VCF along with one of the transcriptomes’ VCF, reporting back the TP, FP and FN values per mean depth (i.e. average coverage of the SNV in all samples in which it was found) and also the specificity, which was calculated as TP/(TP+FP). Our script also produced a new VCF in which each SNV was tagged as either TP, FP or FN for use in downstream analyses.

### Detection of RNA-DNA differences

It was expected that WES data should inherently contain more SNVs due to a greater coverage and range than RNAseq data. We also investigated those SNVs found in RNAseq but not WES. A variety of possible factors influenced calling these FP SNVs and so we removed SNVs called as FP when (i) the FP SNP was found in a region not passing HaplotypeCaller filters in WES, and therefore lacking coverage to be sure the SNV was not found in WES, and (ii) when the SNV was different from the reference in the bovine genome but was found fixed in our data. We refer to the remaining FP SNVs as ‘RNA-DNA differences’ (RDDs), as coined previously (Li *et al.* 2011). To filter RDDs we used methods based on further work arising in the field (Pickrell *et al.* 2012; Kleinman and Majewski 2012; Lin *et al.* 2012). Three relevant methods were used to determine if RDDs were due to sequencing-related factors: (i) RDD was at the extremity of reads supporting it; to resolve this we took all reads in BAM supporting RDDs at the sample level and if all were within 5 bp of either end of the reads, we removed them; (ii) if the RDD was within 5 bp of any splice junction from our database created in STAR, we removed them; (iii) if the RDD was within 5 bp of any other RDD, we removed them both due to potential indel or other variants causing the SNV.

Finally, to address differences between TP and RDD in each transcriptome, we investigated absolute and proportional occurrence of each type of mutation, i.e. G>A or C>G etc. As our transcriptome data was not strand-specific, we could not be certain of the direction of the mutation, and therefore grouped reciprocal mutations i.e G>A and T>C, together. Finally to investigate the properties of RDDs we calculated transition/transversion ratio (Ts/Tv) using total numbers of each reciprocal mutation catagory and tested whether the difference in the Ts/Tv ratios in RDDs was significantly different to that from the TP dataset using a Chi Squared test (p<0.05).

### Functional Annotation of SNVs

The cows sequenced were part of a model of fertility. Therefore, it was of interest to identify SNVs found divergent between the two phenotypes (Fert+ and Fert-). We analysed WES, TPs and RDDs individually for each phenotype. Using snpEff (Cingolani *et al.* 2012) we annotated all SNVs and identified those that were non-synonymous (NS). Due to their potential to cause differences in the structure and functions of a gene, we limited our functional annotation to this set of NS SNVs. We also identified those NS SNVs that were unique to one phenotype or the other in each of WES, TPs and RDDs. Finally, we compared these to the sets of genes found to be differentially expressed in the RNAseq analysis from which the transcriptomes were derived.

